# MegaTrack: a framework for the anatomically accurate and time-efficient virtual dissection and analysis of large-scale tractography datasets

**DOI:** 10.1101/2025.05.27.656234

**Authors:** Flavio Dell’Acqua, Ahmad Beyh, Richard Stones, Rachel LC Barrett, Francisco De Santiago Requejo, Pedro Luque Laguna, Luis Miguel Lacerda, Catherine Davison, Anoushka Leslie, Henrietta Howells, Laura H Goldstein, Steven CR Williams, Marco Catani

## Abstract

The increasing prevalence of large-scale neuroimaging initiatives has created a need for efficient and anatomically accurate methods for analysing tractography data. Here, we present MegaTrack, a framework designed to combine the anatomical precision of manual dissections with the efficiency required for big data analysis. At its core, MegaTrack operates by normalising individual tractography datasets to a standard space, concatenating them into a single “mega” tractography dataset the user manually dissects before automatically extracting subject-specific tracts and metrics in native space. This approach permits identifying and extracting tracts not yet available in existing templates or automatic dissection tools.

We validated this approach using multiple datasets and applications. Comparison with individual manual dissections showed high spatial agreement (weighted dice coefficient > 0.95) and strong correlations in tract-specific metrics (R^2^ > 0.9). In a longitudinal dataset, MegaTrack demonstrated improved reproducibility across time points and higher inter-rater reliability compared to manual dissections. In a motor neuron disease cohort, MegaTrack replicated known group differences in corticospinal tract microstructure with comparable sensitivity to manual dissections (p = 0.001) but with significantly reduced processing time. Finally, we showcase MegaTrack’s scalability by creating a comprehensive white matter atlas from 140 healthy adults and demonstrate its application for lesion analysis in stroke patients. This framework is accompanied by an interactive online tool that allows users to explore these atlases and perform customised lesion analyses.

The MegaTrack framework is a time-efficient solution for the anatomically precise analysis of large tractography datasets, enabling the rapid simultaneous dissection of multiple subjects while preserving individual anatomical features. The framework’s ability to efficiently process large datasets makes it an excellent tool for generating training data for machine learning and deep learning algorithms in tractography analysis. This approach has broad applications in both research and clinical settings, from group comparisons to personalised lesion mapping, and can significantly contribute to the development of advanced automated tractography methods.

## INTRODUCTION

In recent years, neuroscience has moved from individual laboratory-generated research to major multi-centre initiatives like the Human Connectome Project (HCP, [Van Essen et al., 2013]), Adolescent Brain Cognitive Development (ABCD, [Casey et al., 2018]), Alzheimer’s Disease Neuroimaging Initiative (ADNI, [Weiner et al., 2010]), and the UK Biobank (UKBB, [Sudlow et al., 2015]). In the neuroimaging field, this shift has produced unique datasets whose richness is represented by the large number of participants and the complexity and volume of data collected for each. Such data have been used to further our understanding of the anatomy and function of normal and pathological brains. For instance, scientists have proposed new parcellations of the neocortex based on converging data streams from large cohorts [Glasser et al., 2016; Schaefer et al., 2018], gained more profound insights into the role of network interactions in the development of psychosis symptoms [Parkes et al., 2021] and the heterogeneity of grey matter volume changes implicated in multiple psychiatric diagnoses [Segal et al., 2023].

Clearly, big data offers opportunities to investigate the human brain on a large scale but inevitably poses significant challenges that require specific approaches. This is particularly important for tractography, the only available imaging technique to non-invasively reconstruct the major white matter tracts in the living human brain. Tractography has been successfully used to dissect tracts and quantify white matter abnormalities in brain disorders such as autism [Catani et al., 2016], stroke [Forkel et al., 2014], the Riddoch syndrome [Beyh et al., 2024], childhood maltreatment [Lim et al., 2024], and dementia [Amlien and Fjell, 2014], and to describe novel aspects of typical human connectional anatomy [Arrigo et al., 2016; Beyh et al., 2022; Catani et al., 2012; Catani et al., 2017; Thiebaut de Schotten et al., 2011a; Yeatman et al., 2014]. Tractography is widely applied as a quantitative tool for white matter segmentation and represents the basis for structural connectivity analysis [Zalesky et al., 2025]. Compared to voxel-based analyses, tractography has the unique ability to map the anatomy of single tracts and characterise their variability across individuals.

Two tractography approaches have been generally adopted to visualise and extract quantitative measures of white matter. In relatively small datasets, manual dissections performed by expert users for each subject produce highly accurate results comparable to post-mortem dissections [Aarnink et al., 2014; Catani et al., 2002; Tunç et al., 2014; Wassermann et al., 2016]. Additionally, manual dissections have the advantage of adapting the dissecting pipeline to the anatomy of each subject, thus preserving interindividual variability. Nevertheless, manual dissection is particularly laborious and time-consuming, which makes it difficult to extend to large datasets.

Alternatively, automatic dissections allow for the efficient segmentation of white matter tracts in larger datasets using either a predefined template of anatomical landmarks [Yeatman et al., 2012] or clustering approaches based on geometrical descriptors of tract shape [Garyfallidis et al., 2018; Guevara et al., 2012; Wasserthal et al., 2019]. Automatic pipelines permit the fast analysis of large datasets, but they leave little or no room for user interaction to control or check for artefactual results, often yielding lower anatomical precision [Guevara et al., 2012; Lebel et al., 2008]. More recently, initial results from a new generation of automatic tractography dissecting techniques using advanced machine and deep learning strategies are promising [Benou and Riklin Raviv, 2019; Neher et al., 2015; Neher et al., 2017; Wasserthal et al., 2019; Wegmayr et al., 2019]. Nevertheless, they require a significant amount of high-quality training data to accurately learn the anatomical features of each tract and its intrinsic variability across subjects. Common to most existing automatic approaches is the need for predefined templates or anatomical regions for each tract. New tracts cannot be included in automatic pipelines until the corresponding templates or anatomical rules are provided, often through the laborious manual dissection approach.

In this study, we propose MegaTrack, a novel approach for the anatomically accurate and time-efficient dissection and analysis of large tractography datasets. MegaTrack is a framework that enables users to perform simultaneous manual dissections of multiple tractography datasets and to automatically extract tract measurements from each dataset in its native space. Our aim is to develop a framework designed to facilitate tractography data extraction in large cohorts or studies tailored to smaller groups selected for specific demographic, clinical, or neuropsychological characteristics. This framework is not limited to cross-sectional analysis but can also be applied to longitudinal studies and can be used to generate customised atlases of already known or newly discovered white matter pathways; it can also be used for tract lesion analysis and to generate accurate training data for novel machine learning applications. Originally introduced at the 23rd Annual Meeting of the International Society for Magnetic Resonance in Medicine [Dell’Acqua et al., 2015], the MegaTrack framework has since evolved. This paper introduces a major and comprehensive extension of our framework, presenting extensive applications, validation results, and a new publicly available online viewing and analysis tool at https://megatrackatlas.org.

The MegaTrack pipeline consists of four steps as visualised in Figure 1. It begins by remapping each subject’s tractography streamlines to a standard anatomical space and merging them into a single “mega” dataset for manual dissection. This approach enables users to correct artefacts, identify neuroanatomical landmarks in standard space, and dissect streamlines across all subjects simultaneously. By assigning a unique subject and streamline ID (SSID) to each streamline, native-space tractography can be automatically recovered and tract-specific measurements readily extracted. Additionally, identifying common anatomical landmarks in template space facilitates along-tract analysis by allowing precise segmentation of the same tract portions across individuals, thereby improving consistency in quantification and dissection.

**Figure 1.**
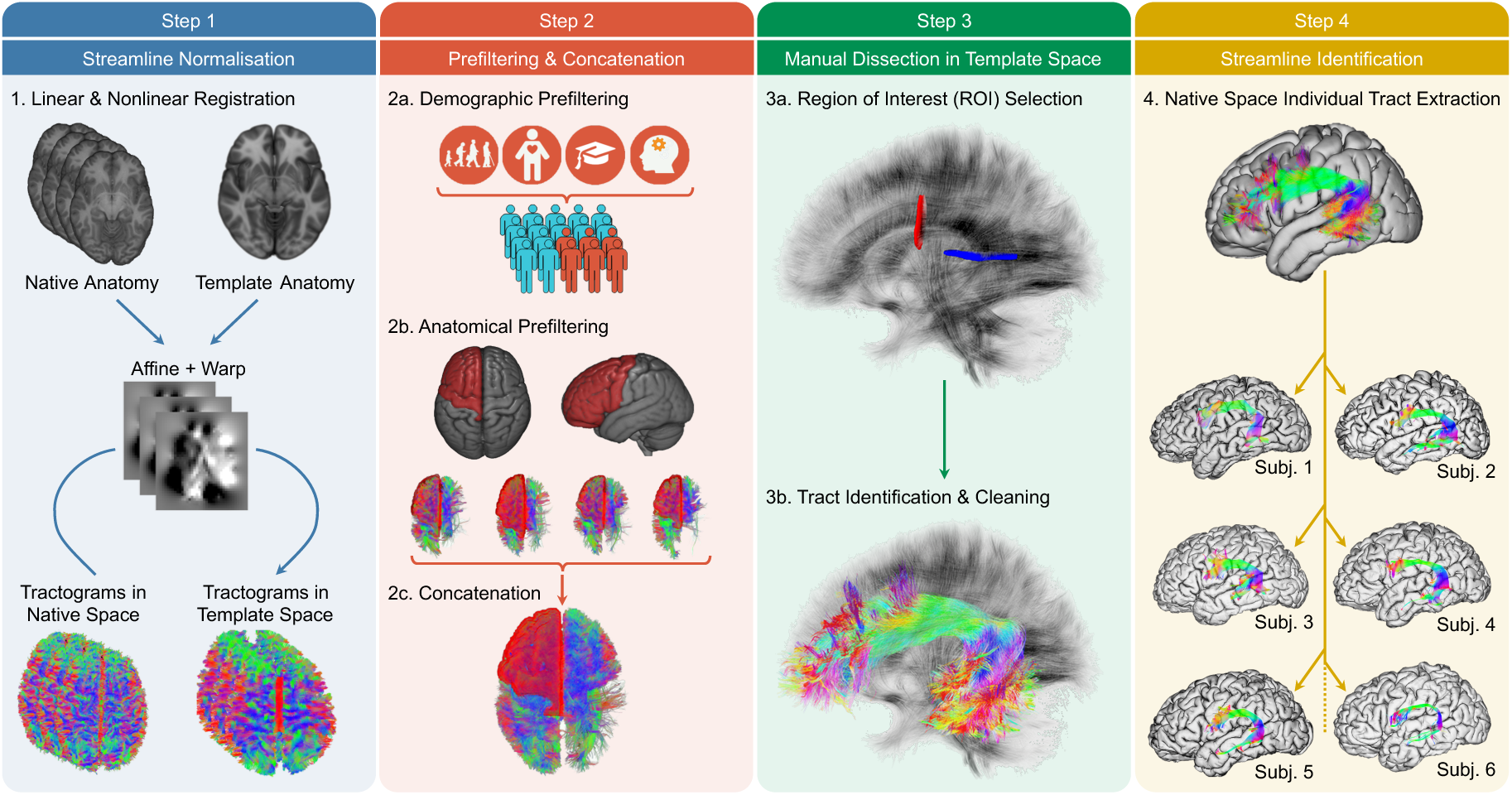
The MegaTrack data generation workflow. This figure illustrates the four main steps of the MegaTrack data generation workflow: (1) streamline normalisation, where individual tractography datasets are mapped to a standard space; (2) prefiltering and concatenation, which involves demographic and anatomical filtering before combining datasets; (3) manual dissections in template space, where tracts are identified and artefacts removed on the combined dataset; and (4) streamline identification, where individual tracts are extracted back in each subject’s native space. This step also includes extracting tract-specific metrics from each tract in native space. Additional analysis steps can be included after step 4, as explained in the text.

Here, we apply the proposed approach to several tractography datasets. First, we use a longitudinal dataset of healthy adult subjects to compare MegaTrack with individual manual dissections and to assess their respective reproducibility and inter-rater reliability scores across time points. We then demonstrate multiple applications of MegaTrack to different healthy and clinical tractography datasets. We apply MegaTrack to a motor neuron disease (MND) dataset to demonstrate its ability to detect group differences of known biological changes and compare it with traditional manual dissections. We also perform dissections of an adult dataset of 140 subjects to generate reference atlases for some of major white matter pathways. Finally, we apply the MegaTrack framework to an example stroke lesion analysis where we automatically assess the probability of white matter tract involvement based on population connectivity data.

## METHODS

### Theory and workflow

The general MegaTrack framework consists of four distinct initial processing steps (**Figure 1**): (1) tractography normalisation to standard space; (2) MegaTrack dataset generation; (3) simultaneous supervised anatomical dissections; and (4) single-subject streamline extraction and analysis. The last step can be performed in native and/or standard space, depending on the analysis required. These steps form the basis of the broader MegaTrack workflow, which supports a wide range of downstream analyses. In the following sections, we explain each stage and provide an overview of the full pipeline.

#### Step 1: Tractography normalisation to standard space

Normalising tractography datasets to a standard space involves two main steps (**Figure 1**, Step 1). First, affine and non-linear transformations are calculated to map each subject’s anatomy to a common standard space (e.g., MNI152). Different registration approaches can be adopted for this initial step, provided that at least one affine transformation and one deformation field are generated. The most straightforward approach is to use image-based registration to directly map each subject’s Fractional Anisotropy (FA) map to an FA template using widely available registration tools such as FSL (https://fsl.fmrib.ox.ac.uk/) or ANTs (https://github.com/ANTsX/ANTs). Advanced tensor-based registrations can also be computed by using, e.g., the TORTOISE software package (https://tortoise.nibib.nih.gov/) or DTI-TK (https://dti-tk.sourceforge.net/). Compared to T1-weighted images, FA and tensor images provide richer information about the anatomical features within white matter and are, therefore, expected to yield a more accurate alignment of deep white matter features [Irfanoglu et al., 2016].

Once all affine transformations and non-linear warping fields have been calculated, normalised tractography datasets can be obtained by applying these transformations and remapping each streamline point to its new position in common space. Importantly, given that the mapping is applied to each streamline point independently, preserving a uniform streamline step size is not required when moving to standard space. Also, compared to traditional scalar image transformation, where intensity interpolation is required to create the final image, no smoothing or loss of information is introduced to streamlines during the operation; the streamlines are simply repositioned and reshaped to match the template’s anatomy.

#### Step 2: MegaTrack dataset generation

A MegaTrack dataset is generated by concatenating the individual normalised tractography datasets obtained from the previous normalisation process into a single “mega” tractography dataset (**Figure 1**, Step 2). During this stage, each streamline is tagged with two ID numbers: a Subject ID linking the streamline to its original subject and a Streamline ID corresponding to the streamline’s index within its native tractogram. This tag, abbreviated as SSID, ensures that every streamline in the MegaTrack tractogram can always be traced back to its native dataset without relying on its relative position within the MegaTrack dataset or inverse spatial transformations.

The concatenation step offers the flexibility to prefilter the MegaTrack dataset based on demographic or anatomical priors. For instance, demographic prefiltering can include only subjects meeting specific criteria such as age, sex, handedness, or IQ, if this information is available. Additionally, anatomical prefiltering can be applied to constrain the MegaTrack dataset to specific brain regions, such as a hemisphere or a single lobe. In practice, we have observed that this prefiltering step can reduce the number of included streamlines by a factor of 10 or more.

Depending on the final number of included streamlines, handling a MegaTrack dataset can require a large amount of memory and computational resources. In this case, further data reduction can be achieved by applying fibre compression to each streamline. This works by reducing the number of points that define a streamline’s spatial trajectory while controlling for the introduced anatomical error [Presseau et al., 2015]. For example, if the error is limited to <0.4 mm, an additional ten-fold compression may be achieved. Again, because the original native streamlines are recovered later by their SSID for the final analysis, no accuracy or information are lost with this fibre compression step.

#### Step 3: Simultaneous supervised dissections

One of the unique features of MegaTrack is the ability to display and dissect streamlines from all subjects within a standard anatomical space (**Figure 1**, Step 3). This offers two key advantages: (1) it enables the user to efficiently and simultaneously perform virtual dissections across multiple subjects while (2) maintaining unbiasedness to the individual anatomical variability. MegaTrack allows an expert operator to perform traditional ‘manual dissections’ using manually defined waypoint regions of interest (ROIs), atlas-based subcortical ROIs, cortical ROIs (e.g., obtained from FreeSurfer), or a combination of these three approaches. Geometrical constraints such as length or shape filtering may be easily applied to select certain populations of white matter tracts, e.g., U-shaped fibres [O’Halloran et al., 2017]. Throughout the dissection process, the operator can identify and remove artefactual streamlines much like they would during single-subject manual dissections. The interactive nature of these dissections highly benefits from the group-level approach, as the operator is continuously guided by the average anatomy of the underlying template and, at the same time, by the outliers and artefactual streamlines of individual subjects. This approach also minimises unintended errors in ROI placement that might occur during manual ROI delineation in individual datasets, since the same anatomical landmarks are now used for all subjects.

#### Step 4: Individual streamlines extraction from MegaTrack to subject space

The final step extracts each subject’s dissected streamlines from their native tractogram (**Figure 1**, Step 4). Once MegaTrack ROIs are defined, reading the SSIDs from the dissected MegaTrack streamlines allows for the direct extraction of individual dissected tracts in native space. This is achieved by simply selecting the desired streamlines in each subject’s native tractogram, without the need for spatial transformations. The analysis and a list of possible applications of the MegaTrack framework resulting from this step are covered in the next section.

### Analysis and applications

Following extraction of individual tracts for each subject, the MegaTrack framework offers several analysis options and tractography applications. In this study, we first assess the reproducibility and reliability of tract-specific measurements obtained using MegaTrack against individual manual dissections. We then investigate three potential applications: case-control group comparison, atlas generation, and lesion mapping.

We used four datasets, each processed using different parameters commonly chosen in the neuroimaging research literature, using both DTI and spherical deconvolution (SD) tractography. This allowed us to assess the MegaTrack’s utility across different acquisition and processing settings. Full details on each dataset, including acquisition and processing parameters, are available in the Supplementary Material.

#### Application 1: Reproducibility and reliability of tract-specific measurements

*<u>Motivation:</u>* MegaTrack offers automated extraction of tract-specific measurements for large studies, potentially providing more uniform and reproducible dissections across subjects, time points, and users compared to manual methods. By dissecting all subjects simultaneously on a common template, MegaTrack aims to reduce variability associated with manual dissection over time or between users (training effects, user fatigue, consistency, etc). Once tracts and metrics are extracted, standard tract-specific and along-tract analyses can be performed.

*<u>Analysis:</u>* To test this, we compared MegaTrack and individual manual dissections using a longitudinal dataset from 16 healthy adults (three sessions per subject within a month). Diffusion data were acquired along 60 diffusion-weighted (DW) directions with a b-value of 1500 s/mm^2^ and an isotropic voxel (2×2×2 mm^3^) (see Dataset 1 in Supplementary Material). We performed DTI-based tractography dissections of the left long segment of the arcuate fasciculus, and spherical deconvolution-based dissections of the left cingulum. First, we assessed the agreement between manual and MegaTrack dissections using the weighted-dice coefficient [Cousineau et al., 2017] on tract density maps and by correlation analysis between the corresponding diffusion and tractography metrics. Second, we evaluated the reproducibility of the diffusion metrics extracted from the cingulum across three time points and three users for both manual and MegaTrack dissections using the Intraclass Correlation Coefficient (ICC), specifically the ICC(2,1) type, as described in Shrout and Fleiss [1979]. Third, we calculated the inter-rater reliability across the three users, considering all time points using again ICC(2,1).

#### Application 2: MegaTrack for group comparison analysis

*<u>Motivation:</u>* By enabling the simultaneous dissection of multiple subjects, the MegaTrack framework naturally facilitates rapid group-level and longitudinal tractography analyses. We aimed to determine if tract-specific and along-tract metrics obtained from MegaTrack could provide comparable results with similar sensitivity to group differences as those obtained from individual manual dissections.

*<u>Analysis</u>*: To this end, we analysed a dataset of 25 limb-onset motor neuron disease (MND) patients and 24 age- and sex-matched controls, comparing the detection of group differences in the left corticospinal tract using our method against manual dissection results. We acquired this dataset along 32 DW directions with a b-value of 1350 s/mm^2^ and 2.4×2.4×2.4 mm^3^ isotropic voxels (see Dataset 2 in Supplementary Material). We used spherical deconvolution tractography and extracted the hindrance modulated orientational anisotropy (HMOA) index as a proxy measure of fibre density [Dell’Acqua et al., 2013]. Group comparisons were conducted with a two-sample t-test, and Spearman rank correlations between HMOA values and the clinical scores of limb and bulbar symptom severity assessed using the revised Amyotrophic Lateral Sclerosis Functional Rating Scale (ALSFRS-R) [Cedarbaum et al., 1999].

#### Application 3: Scalability and atlas generation

*<u>Motivation:</u>* Creating atlases that represent common brain anatomy and inter-individual variability is paramount but challenging. The proposed framework enables rapid generation of anatomical tracts from multiple subjects in native and standard spaces, leveraging existing deformation fields within MegaTrack. This facilitates direct creation of tract probability maps and population-level average volume and variance estimates. Customised normative atlases can then be generated based on demographics or neurobehavioral characteristics, allowing for further data mining.

*<u>Analysis</u>:* First, to assess MegaTrack scalability and time efficiency, we computed the time required for tract dissection (including all processing steps) using manual method versus MegaTrack approach, varying the number of datasets and tracts. We assumed 15 minutes per single manual tract dissection and 90 minutes for MegaTrack, with a maximum of 40 hours per week. Second, to demonstrate atlas feasibility, we dissected 20 major association tracts per hemisphere and commissural tracts in a HARDI dataset (b=3000 s/mm^2^, 60 DW directions, 2.4×2.4×2.4 mm^3^ voxels) from 140 healthy adults (age 18-80), processed with DTI and SD tractography (see Dataset 3 in Supplementary Material). This amounted to over 5000 dissected tracts. We then generated tract probability maps by averaging the binary masks of the individual tract density maps across all subjects. These dissections are publicly available for visualisation and demographic filtering through the online MegaTrack tool.

#### Application 4: Novel lesion mapping and disconnection analysis

*<u>Motivation:</u>* The MegaTrack framework can introduce a novel approach to lesion analysis by dynamically generating white matter atlases customised to individual patient characteristics (age, sex, etc.). Hence, MegaTrack can be used to generate a customised group dataset and assess the involvement of white matter pathways in single or group patient studies. Using a normalised lesion mask from any structural MRI (or even CT), tracts intersecting the lesion can be identified on the customised atlas, and their disconnection degree quantified with various metrics.

*<u>Analysis:</u>* We demonstrated this approach by analysing a representative lesion mask from a stroke patient with aphasia using a reference atlas generated with MegaTrack. We obtained the mask from a separate imaging study in MNI space. We then calculated three metrics of lesion-tract intersection: (1) *overlap* score, (2) *maximum probability* score, and (3) *disconnection* score. We define the *overlap score* as the voxel fraction of the binarised tract probability maps overlapping the lesion mask. We define the *maximum probability* score as the value of the voxel with the highest probability from a given tract of the atlas overlapping the lesion [Thiebaut de Schotten et al., 2011b]. Finally, the *disconnection score* is a novel lesion-streamline disconnection measure that we obtain by averaging, across all the subjects of the atlas, the percentage of streamlines intersected by the lesion in each subject—that is, the fraction or percentage of streamlines potentially “disconnected” by the lesion. This tract lesion analysis is integrated into the online tool.

## RESULTS

### Comparison with individual manual dissections

Visual comparison (**Figure 2**) of the long segment of the arcuate fasciculus (DTI-based, panel A) and the cingulum (SD-based, panel B) revealed high similarity between manual dissection and MegaTrack approaches, indicating comparable preservation of individual anatomical variability. Statistically, weighted-dice coefficient analyses confirmed excellent spatial agreement for both methods (arcuate DTI, w-dice = 0.99 ± 0.01; cingulum SD, w-dice = 0.99 ± 0.01). Furthermore, strong correlations were observed between MegaTrack and manual dissections (**Figure 2**) for both macrostructural measures (streamline count: arcuate DTI, R^2^ = 0.97; cingulum SD, R^2^ = 0.91), and microstructural diffusion metrics (arcuate FA, R^2^ = 0.99; cingulum HMOA, R^2^ = 0.98). These results confirm that MegaTrack and manual dissection yield nearly identical set of streamlines and tract-specific measurements from each subject.

**Figure 2.**
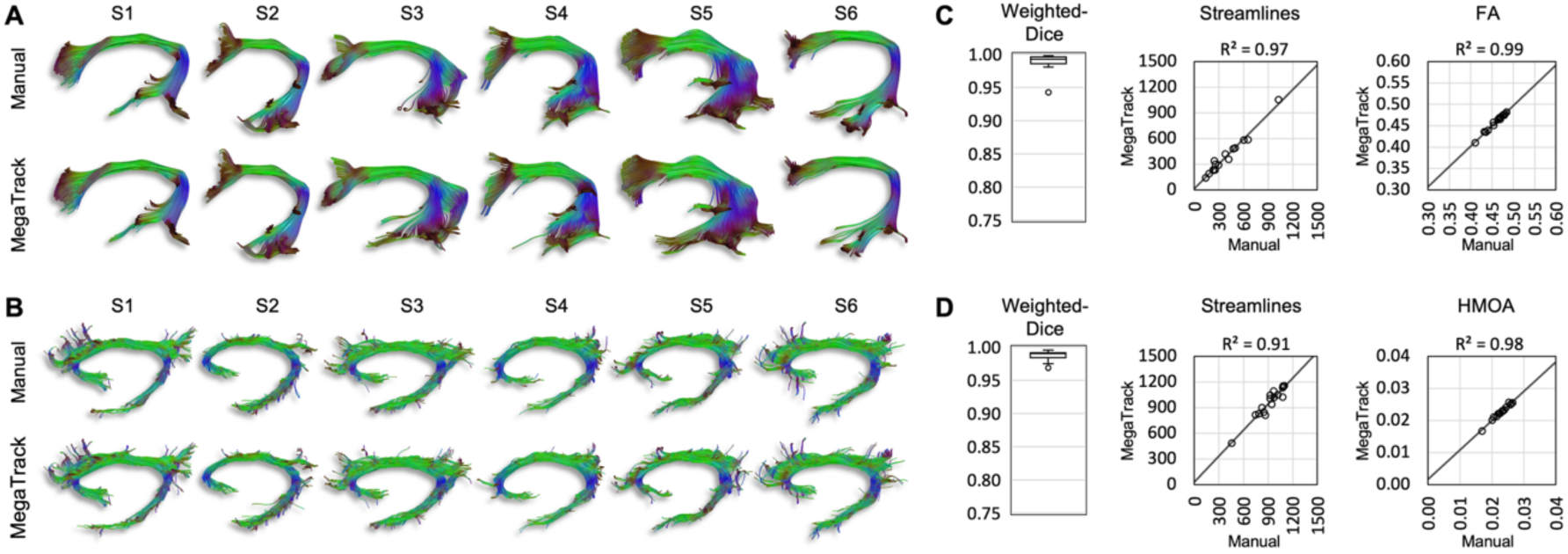
Comparison between MegaTrack and individual manual dissections. The dissections of (**A**) the long segment of the left arcuate fasciculus (DTI) and (**B**) the left cingulum (spherical deconvolution) are shown for six subjects using the individual manual approach and MegaTrack. Panels (**C**) and (**D**) show the correlation graphs between the two approaches using two metrics — the number of streamlines of the final dissected tracts and a microstructural measure (FA for DTI, HMOA for SD) — and the weighted dice coefficient which assesses spatial agreement. Both visual and numerical comparisons indicate a very high level of agreement between the two dissection approaches.

### Reproducibility and inter-rater reliability

To assess reproducibility across time and inter-rater reliability (IRR), three raters, performed manual and MegaTrack dissections of the left cingulum using SD in 16 subjects, each collected at three time points. Raters agreed on dissection criteria before performing dissections. Examples of these dissections from three subjects are shown in **Figure 3**. By visual inspection of dissections from the three time points, all raters showed good reproducibility of the dissected tracts. The difference across raters was minor and clearly smaller than the intrinsic anatomical variability across time and subjects.

**Figure 3.**
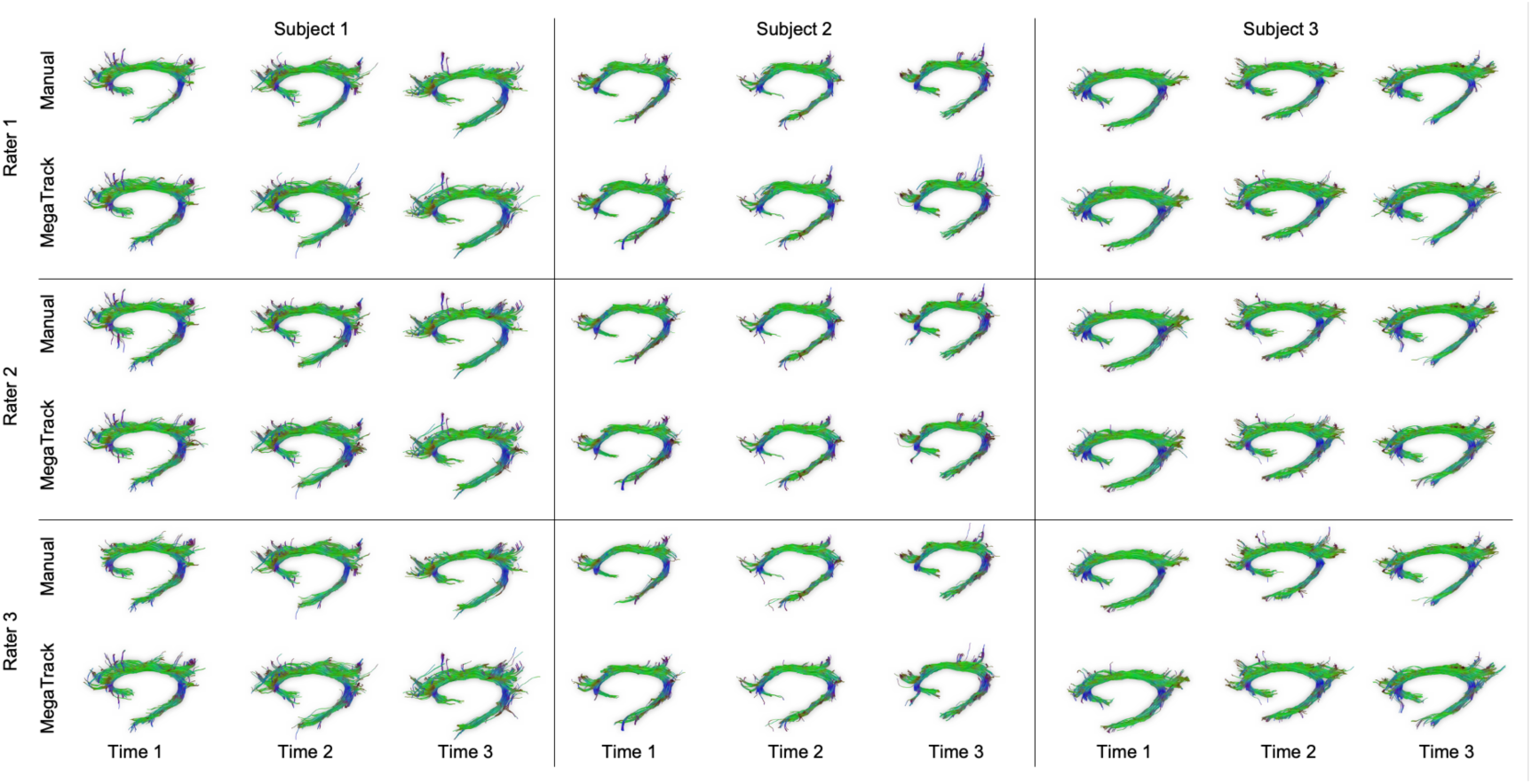
Examples of manual and MegaTrack dissections. Examples of individual manual dissections vs. MegaTrack of the left cingulum (spherical deconvolution) performed in three healthy subjects by three dissectors (raters) for each of the three time points.

Quantitatively, reproducibility measured across the three time points (**Table 1**, left) and inter-rater reliability (**Table 1**, right) were high for both manual and MegaTrack dissections for all measures (number of streamlines, tract volume, FA, and HMOA). Notably, MegaTrack consistently exhibited slightly higher reproducibility across time points compared to manual dissections, suggesting its potential to increase reliability in longitudinal studies. Similarly, MegaTrack showed higher IRR compared to manual dissections between raters indicating reduced variability attributable to rater bias.

**Table 1.** Reproducibility and inter-rater reliability of manual and MegaTrack dissections. Left: reproducibility comparison between manual dissections and the MegaTrack framework defined as ICC computed across three time points. Right: inter-rater reliability comparison between manual dissections and the MegaTrack framework defined as ICC computed across three raters.

|  | ICC Across time |  |  |  |  |  | ICC Across Raters |  |  |  |  |  |
| --- | --- | --- | --- | --- | --- | --- | --- | --- | --- | --- | --- | --- |
|  | Manual |  |  | MegaTrack |  |  | Manual |  |  | MegaTrack |  |  |
|  | ICC | 95% CI Lb | 95% CI Ub | ICC | 95% CI Lb | 95% CI Ub | ICC | 95% CI Lb | 95% CI Ub | ICC | 95% CI Lb | 95% CI Ub |
| # Streamlines | **0.822** | 0.730 | 0.890 | **0.839** | 0.756 | 0.900 | **0.801** | 0.543 | 0.905 | **0.879** | 0.808 | 0.927 |
| Tract vol [ml] | **0.851** | 0.774 | 0.908 | **0.869** | 0.800 | 0.920 | **0.879** | 0.711 | 0.942 | **0.909** | 0.857 | 0.945 |
| FA | **0.779** | 0.673 | 0.860 | **0.793** | 0.692 | 0.869 | **0.919** | 0.847 | 0.956 | **0.972** | 0.955 | 0.983 |
| HMOA | **0.916** | 0.869 | 0.982 | **0.932** | 0.894 | 0.986 | **0.968** | 0.942 | 0.982 | **0.964** | 0.942 | 0.978 |

### MegaTrack for case-control tractography analysis

To evaluate MegaTrack’s suitability for studying microstructural changes in MND, given prior reports of corticospinal tract alterations [Bastin et al., 2013; Nelles et al., 2008; Sarro et al., 2011; Van Der Graaff et al., 2011] we dissected the limb-projecting CST in a group of limb-onset MND patients and matched controls using MegaTrack and manual methods.

MegaTrack-derived HMOA scores revealed a statistically significant difference (*t*(47) = 3.38, *p* = 0.001) between MND patients (0.0209 ± 0.0022) and matched controls (0.0231 ± 0.0023), consistent with the results from the individual manual dissections (patients: 0.0207 ± 0.0024; controls: 0.0230 ± 0.0024; *t*(47) = 3.36, *p* = 0.002). In the patient group, HMOA correlated with the ALSFRS-limb score for both MegaTrack (*r_S_* = 0.44, *p* = 0.019) and individual manual dissections (*r_S_* = 0.48, *p* = 0.012). This effect was anatomically specific as revealed by the lack of correlation between HMOA and the ALSFRS-bulbar score for both MegaTrack (*r_S_* = −0.06, *p* = 0.396, n.s.) and individual manual dissections (*r_S_* = −0.01, *p* = 0.480, n.s.).

### Scalability

The scalability analysis for manual and MegaTrack dissections is reported in **Figure 4**. For small studies (20 subjects, 2 tracts) both methods required a similar time investment of approximately two to three working days, with MegaTrack’s initial normalisation and concatenation process of all datasets matching the time needed for manual dissection. Scaling the analysis to include more tracts or subjects resulted in a linear increase in processing time for MegaTrack, but a non-linear increase for individual manual dissections. For a large scale study (200 subjects and 10 tracts), the manual method was projected to demand approximately 500 work hours (roughly 13 weeks), whereas the same task could be completed with MegaTrack in just over one week, underscoring its efficiency for larger datasets.

**Figure 4.**
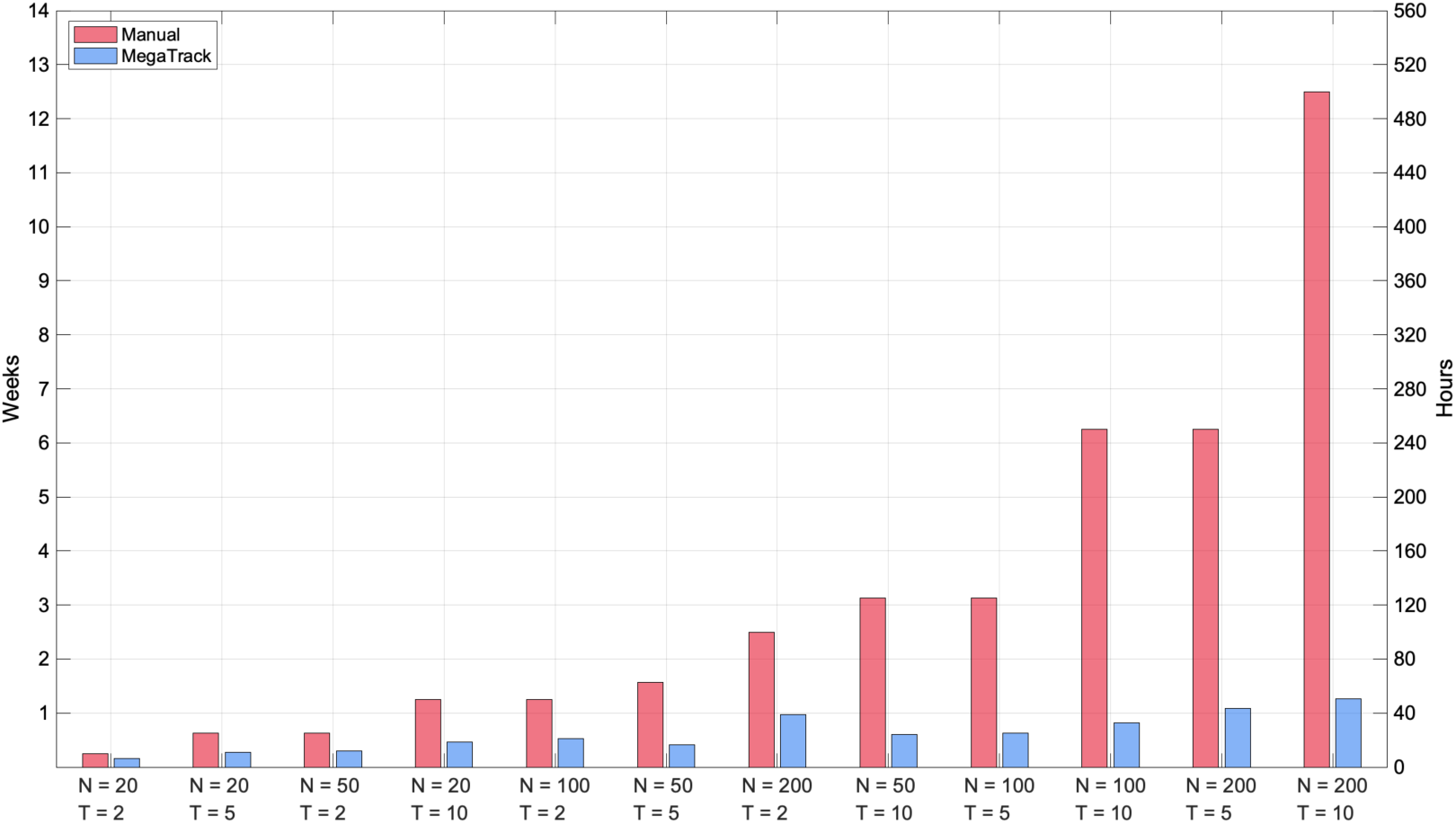
MegaTrack scalability analysis. This plot depicts the simulated time required for a project based on various combinations of numbers of participants and dissected tracts, for both the individual manual dissection approach (red) and the MegaTrack approach (blue). Each point along the x-axis corresponds to a combination of sample size (N) and number of dissected tracts (T). The left-hand side y-axis depicts time as the number of calendar weeks required to complete the project, which is based on an eight-hour workday and a five-day work week, while the right-hand side y-axis depicts time as the number of hours required to complete the dissections. For example, a study that requires the dissection of T = 10 tracts in N = 200 subjects can take close to 13 weeks (or 500 hours of work) to complete following the manual approach but would be concluded in just over a week (or 50 hours) using MegaTrack.

### Atlas generation

The scalability of MegaTrack facilitates the rapid generation of anatomically vetted, large-scale atlases of multiple white matter tracts. To demonstrate this, we have used a dataset of 140 healthy subjects (age 18-80) to create DTI and SD tractography atlases of the major associative and commissural white matter tracts. Probabilistic maps for the cingulum and fornix reconstructions based on SD tractography from the complete cohort and displayed on the MNI152 template are shown in **Figure 5**. An interactive version of these atlases is publicly available through the MegaTrack website for a comprehensive exploration of all tracts. An important feature of this application is the ability to select specific cohorts by filtering the tractography results based on demographic characteristics. The website enables the user to generate customised tractography atlases tailored to specific age groups, sexes, or other population-specific characteristics, enhancing the precise and relevance of analyses in research and clinical applications.

**Figure 5.**
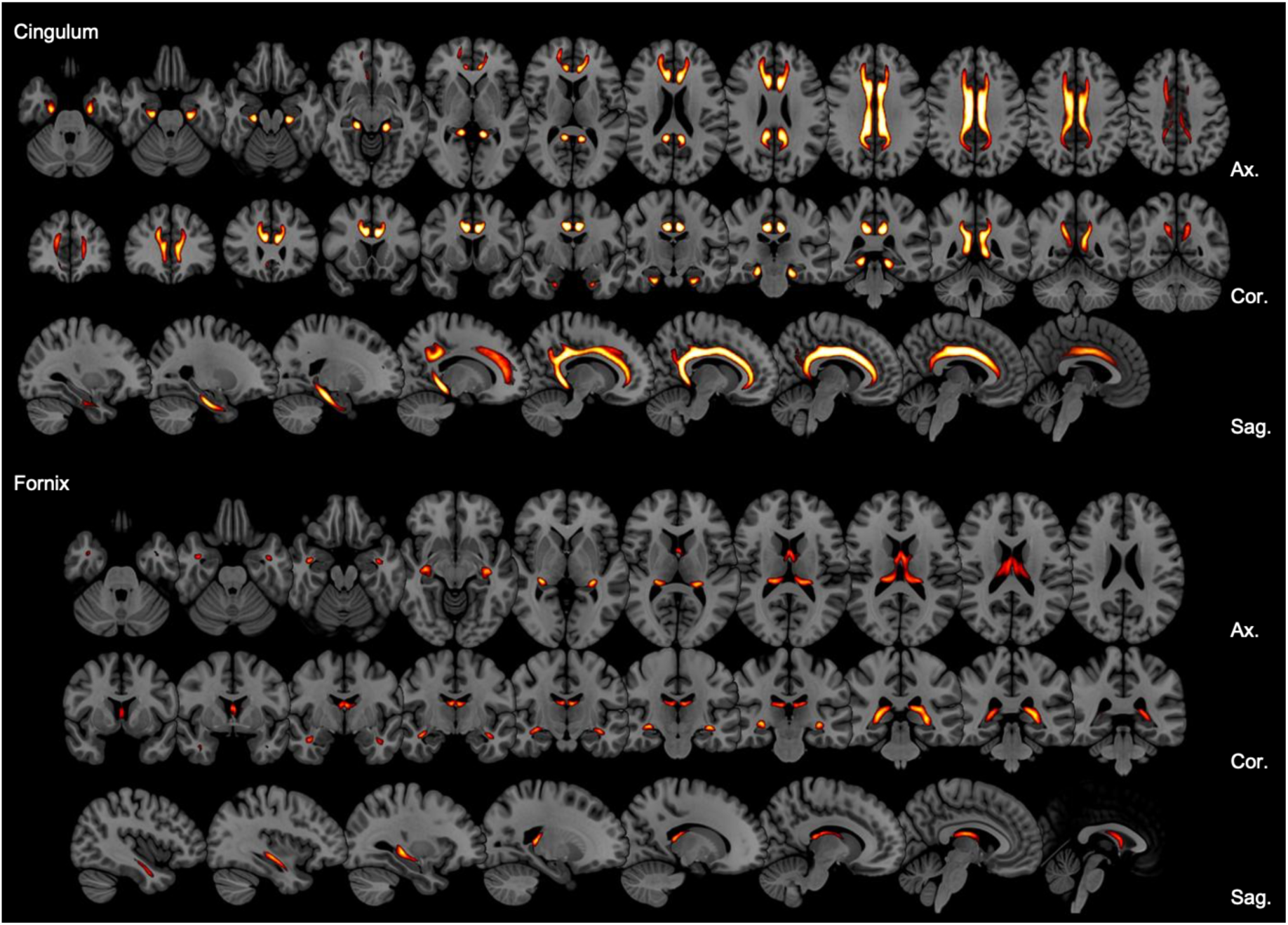
MegaTrack atlases. Examples of two probabilistic atlases obtained directly from the MegaTrack framework after dissecting the dataset of 140 healthy subjects. In the top panel, the left and right cingulum are displayed through the most representative axial, coronal, and sagittal slices in MNI space. The bottom panel shows the same visualisation for the fornix.

### Lesion mapping using MegaTrack-derived atlases

When applied to clinical populations, MegaTrack can enhance lesion mapping approaches and their clinical-anatomical correlation inferences. As a proof of concept, **Figure 6** shows the results of two widely used lesion mapping approaches (A and B) and a novel approach based on MegaTrack (C) applied to a stroke patient with aphasia, a syndrome known to be associated with a disconnection of the language network.

**Figure 6.**
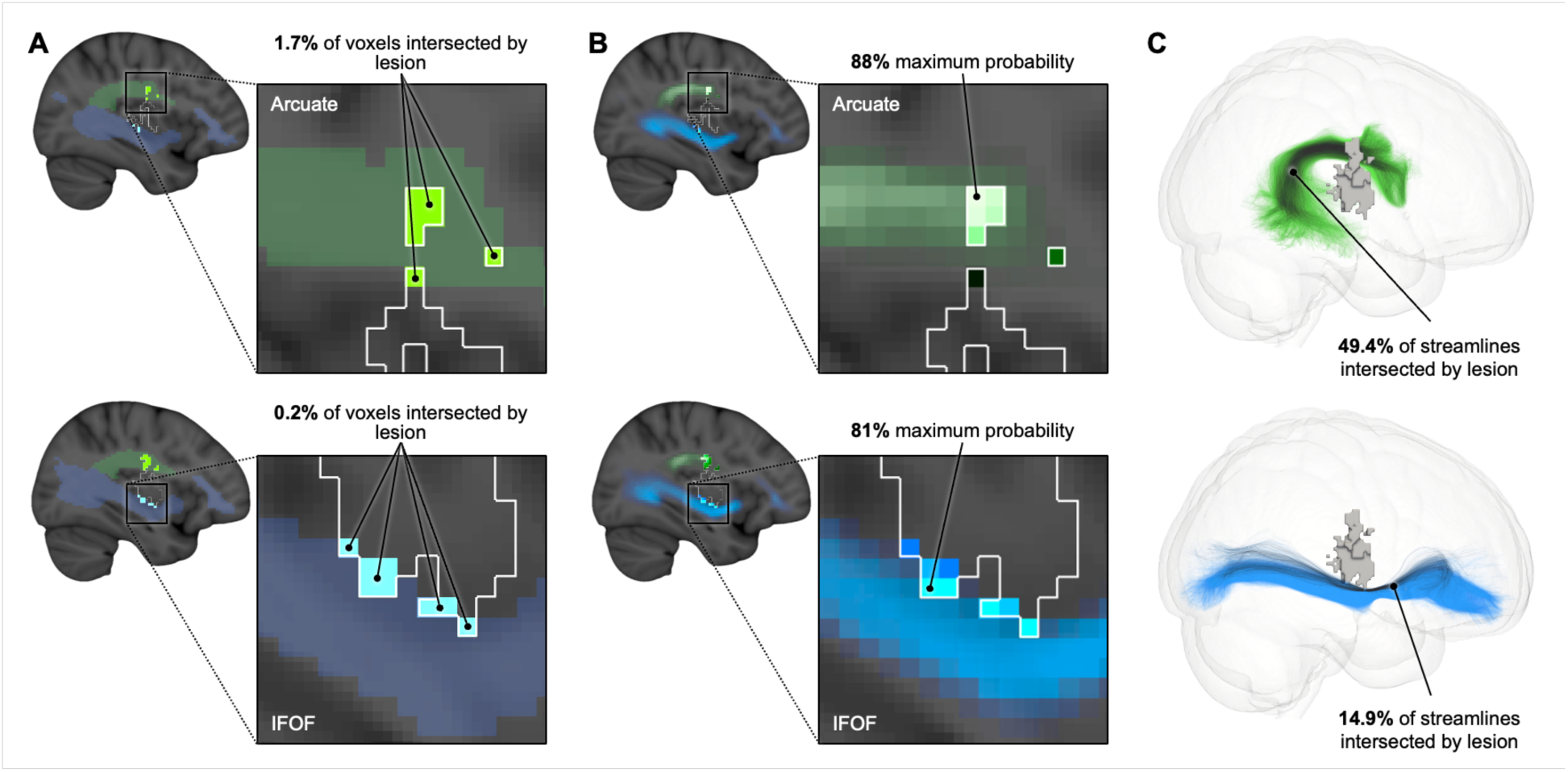
Lesion mapping using three different approaches. Three approaches are used to assess the potential impact of a stroke lesion on two tracts, the arcuate fasciculus (AF) and the inferior fronto-occipital fasciculus (IFOF). (**A**) The lesion-voxel overlap approach, which calculates the fraction of tract voxels intersected by the lesion, shows a marginal damage (<2%) to both tracts. (**B**) The lesion-max probability analysis, which extracts the highest tract-lesion intersection probability (subject overlap score), shows a high probability of involvement (>80%) of both tracts. (**C**) The lesion-streamline approach, which calculates the fraction of streamlines intersected by the lesion, identifies a higher degree of disconnection for the arcuate fasciculus (approx. 50%) compared to a much lower involvement of the IFOF (approx. 15%).

While all three lesion mapping methods identified damage to both tracts in this patient, the results exhibited significant differences. The lesion-voxel overlap approach (Figure 6, A) utilises binary maps derived from a reference group to identify the number of voxels composing a tract map intersected by the lesion. In the aphasia patient, by thresholding the tract probability maps at a minimum of 25% (i.e. at least 25% of subjects have streamlines passing through each voxel of the tract map), the corresponding lesion-tract voxel overlap percentage was 1.7% for the AF and 0.2% for the FOF, suggesting a limited involvement of these tracts.

The method of maximum probability utilises probability maps in which a number is assigned to each voxel composing a tract map and that number corresponds to the percentage of subjects in the reference group whose tract occupies the voxel. The analysis indicates the voxel with highest probability intersected by the lesion, which in this case was very high for both the AF and IFOF at 88% and 81%, respectively. These results indicate that across the whole population defined by the atlas there is a high probability that both tracts are affected by the lesion. However, these two analyses do not provide a complete measure of the degree of disconnection nor a clear indication of which of the two tracts may be more involved in the clinical symptoms.

In contrast, the MegaTrack-based lesion mapping method offers a more direct measure of streamline disconnection across the affected population. This method maps the lesion directly on the tractogram generated from the reference group, thus avoiding the biases linked to the choice of thresholds for the creation of the binary masks or the overestimations derived from the probability maps. The disconnection scores for the AF and IFOF, were 49.4% and 14.9%, respectively, indicating a more considerable involvement of the AF, which aligns well with the aphasia symptoms displayed by patients with this type of lesion.

## DISCUSSION

The neuroimaging field today is increasingly reliant on the use of large datasets acquired by national and multinational consortia (e.g., HCP, UKBB, EU-AIMS). These datasets pose analysis challenges for tractography, which usually relies on approaches carried out in each subject’s native space. This is especially important because the white matter anatomy, i.e., the shape and size of different white matter bundles, is highly variable across individuals [Catani et al., 2007; Croxson et al., 2018; Rojkova et al., 2016; Thiebaut de Schotten et al., 2011b]. So far, in the past the typical tractography approach has been to guarantee anatomical accuracy by performing dissections in native space at the expense of time. This leads us to an important question: how can we analyse large tractography datasets efficiently while maintaining a high level of anatomical accuracy?

A solution is to adopt fully automated tractography dissection methods. These approaches are the fastest but leave little room for user intervention and the final dissections are usually based on hard-coded anatomical priors (regions of interest or tract templates for each white matter tract). Similarly, emerging machine learning methods for automatic dissection show promise but require extensive training datasets of high-quality, pre-dissected tracts from expert anatomists. Though these methods continue to improve, they currently offer a limited set of predefined tracts that can be dissected and analysed while, at same time, white matter anatomy and its nomenclature is still rapidly evolving. Being able to customise tractography analysis around newly defined or less known tracts can be essential in certain studies.

Our proposed method, the MegaTrack framework, offers a complementary approach by combining the scalability of automatic dissections with the anatomical precision of manual dissections. MegaTrack starts by non-linearly mapping the streamlines of each subject to a common template to reduce inter-individual variability while preserving the individual anatomical connectivity information carried by each streamline. This enables simultaneous manual dissection of all subjects, guided by the unbiased anatomy of the chosen template, which guarantees the consistent definition of tract boundaries and waypoints across subjects. The framework maintains fine anatomical control through direct user interaction during the dissection process, without being limited by predefined tract templates. Using a unique subject and streamline ID system, MegaTrack enables two-way mapping between the native and template space and allows the instant extraction of corresponding tracts in each individual’s native space along with associated microstructural measures, fully preserving individual anatomy without any loss of information.

Importantly, automatic approaches and MegaTrack can work synergistically. For example, automatic methods can be used to prefilter tractograms before MegaTrack dissection, while vice versa MegaTrack can help validate or refine automatically generated results. Our analysis demonstrated that MegaTrack dissections achieve near-perfect spatial agreement with individual manual dissections and comparable macro- and micro-structural properties. Moreover, our data shows that performing dissections on a standard template decreases dissection variability in longitudinal studies and achieves a better agreement between raters. This gives MegaTrack an advantage over individual manual dissections, because manual dissections can be challenging to perform consistently across individuals or time points due to the changing anatomy of each dataset and the reliability of the user over time. Although we only computed ICC indices across 16 subjects in this study, we expect to see even larger improvements over manual dissections when considering much larger datasets.

To further test the performance of the MegaTrack framework, we applied it to a clinical study of motor neuron disease (MND) and conducted a comparison between individuals with MND and a healthy control group. Both the MegaTrack and manual approaches were successful in revealing the expected statistically significant group difference in the microstructural properties of the cortico-spinal tract, as measured by HMOA. Further, both approaches revealed an anatomically specific relationship between microstructure and clinical scores, captured by a positive correlation between HMOA and the ASLFRS-limb score, and the absence of a correlation with the ALSFRS-bulbar score. In addition, all results from both the MegaTrack and individual manual approaches were highly similar, but with MegaTrack requiring less than an hour for tract dissection and value extraction. In comparison, the individual manual dissections had to be conducted over several days to ensure that each dissection received the required level of rigour. Although we only assessed microstructure using tract-average HMOA, different types of tractography analysis can also be combined with the MegaTrack framework, including more advanced ones like along-tract analysis, or even looking at individual tract segments at selected anatomical positions [Colby et al., 2012; Yeatman et al., 2012].

To further illustrate MegaTrack’s scalability, we estimated the dissection time required for both MegaTrack and individual manual dissections while accounting for factors such as the number of work hours in a day. Our analysis showed that even small studies using data from 20 subjects and dissecting more than two tracts would benefit from using MegaTrack. As expected, the time difference between the two methods increased substantially when we considered a larger number of subjects or a larger number of tracts. In the case of a larger study with 200 subjects and 10 tracts, MegaTrack would still allow the user to obtain results in around a week whereas individual manual dissections would require close to three months of continuous dissections. Additionally, MegaTrack’s main time constraint is not associated with the dissection phase but more with the initial registration and dataset preparation. The dissection of additional tracts only introduces a small-time increment in the context of the whole analysis. Thus, MegaTrack has a clear time advantage over individual manual dissections, which is important not only because it allows for precise and rapid extraction of tracts and metrics, but also because it opens the door, for the first time, to revisions or corrections of entire datasets simultaneously without paying a large time penalty, and without introducing errors into individual dissections.

Combined with its anatomical precision, MegaTrack’s scalability makes it an ideal tool for generating white matter atlases made of a large number of tracts. Given the increasing reliance of the newer generation of machine learning dissection tools on atlases and training datasets, MegaTrack represents an efficient solution for generating training datasets. Further, atlas generation based on specific demographic details allows for a more precise characterisation and stratification of subgroups within larger populations. In this context, MegaTrack can be used to build atlases of white matter anatomy for various age groups and clinical or non-clinical populations, and these can be used to study developmental trajectories through normative modelling [Marquand et al., 2016].

The importance of such atlases becomes clearer when we consider clinical case studies. Here, the power of big data can be harnessed by MegaTrack for single-case clinical applications even when diffusion data is not available for the patient. Based on a clinical structural MRI or CT scan, a user can delineate lesions caused, for example, by stroke, white matter hyperintensities in dementia, or demyelination in multiple sclerosis and analyse the impact of that lesion on white matter networks based on a demographically filtered atlas. Furthermore, in this study we propose a new method to quantify the degree of disconnection of a tract that is facilitated by the MegaTrack approach. This method not only considers the localisation and volume of the lesion but also quantifies the percentage of affected streamlines touching the lesion in each subject of the reference dataset as a proxy of white matter disconnection. Such a tool would be of great value for the wider research and clinical communities, both as an investigative and teaching instrument. Our proposed method offers an additional dimension to better quantify and correlate disconnections with clinical scores. In this spirit, we created an online tool to visualise these atlases and to run lesion analyses.

As with most neuroimaging tools, MegaTrack has limitations that must be addressed. Firstly, because MegaTrack relies on spatial normalisation, the final dissections will be affected by the quality of this normalisation. Here, there is room for improvement by using advanced registration tools tailored for white matter, such as tensor-based spatial normalisation. This also means that using the MegaTrack lesion analysis framework must be restricted to cases for which good spatial normalisation can be achieved. Large abnormalities that considerably deform the brain’s anatomy (e.g., tumours or cysts) may be problematic in this context. Secondly, dissecting very small tracts, such as some U-shaped tracts, or tracts with very large inter-individual variability may be more challenging compared to larger or less variable tracts. Nevertheless, in such cases, the MegaTrack framework can still be used in combination with pre-cleaning or clustering tools, or anatomical and geometric constraints [de Santiago Requejo et al., 2017].

The MegaTrack framework can be further expanded in multiple directions. A main point of improvement is making MegaTrack more computationally efficient and capable of handling even larger datasets made up of thousands of brains. A second key aspect is the integration with additional tractography analysis tools that can benefit from the availability of precise anatomical dissection. These include along-tract analysis, segmenting tracts within selected anatomical references, among others. A third avenue for improvement concerns the use of pre-cleaning methods (e.g., COMMIT, SIFT, iLife) to prefilter the streamlines that enter into MegaTrack, thereby making the disconnection scores of the lesion analysis more closely related to the underlying anatomical fibre density. Finally, the MegaTrack framework can be easily translated to animal models, including rodents and non-human primates, to further accelerate the analysis of tractography data where automatic methods are currently lacking in both anatomical tract templates and anatomical priors.

## ACKNOWLEDGMENTS

FDA was supported by the Catalyst: International Leader Fellowship, funded by the New Zealand Ministry of Business, Innovation and Employment (MBIE) and administered by the Royal Society Te Apārangi.

The MND study was supported by the Wellcome Trust (Grant reference 083477/Z/07/Z). For the purpose of open access, the author has applied a CC BY public copyright licence to any Author Accepted Manuscript version arising from this submission.

This paper represents independent research part-funded by the NIHR Maudsley Biomedical Research Centre at South London and Maudsley NHS Foundation Trust and King’s College London. The views expressed are those of the authors and not necessarily those of the NHS, the NIHR or the Department of Health and Social Care.

The authors would like to thank the members of the NatBrainLab for their support and feedback.

## CODE AVAILABILITY

The MegaTrack code will be made publicly available upon completion of the peer-review process.

## DATA AVAILABILITY

The data generated by this study, the interactive atlas, and the lesion mapping tool, are available through the MegaTrack website: https://megatrackatlas.org.

