## Supplementary Material for "MegaTrack: a framework for the anatomically accurate and time-efficient virtual dissection and analysis of large-scale tractography datasets"

Flavio Dell'Acqua [1,2*], Ahmad Beyh [1,3,*], Richard Stones[1,2], Rachel LC Barrett[1], Francisco De Santiago Requejo[1], Pedro Luque Laguna[1], Luis Miguel Lacerda[1], Catherine Davison[1], Anoushka Leslie[1], Henrietta Howells[1], Laura H Goldstein[4], Steven CR Williams[2], Marco Catani[5, 6]

1. *NatBrainLab, Institute of Psychiatry, Psychology & Neuroscience, King's College London, London, SE5 8AF, UK*
2. *Department of Neuroimaging, Institute of Psychiatry, Psychology & Neuroscience, King's College London, London, SE5 8AF, UK*
3. *Department of Psychiatry, Brain Health Institute, Rutgers University, Piscataway, NJ, 08854, USA*
4. *Department of Psychology, Institute of Psychiatry, Psychology & Neuroscience, King's College London, London, SE5 8AF, UK*
5. *Department of Neuroscience, Imaging, and Clinical Sciences (DNISC) and Institute for Advanced Biomedical Technologies, University "G.D'Annunzio", Chieti-Pescara, Italy*
6. *IRCCS Neuromed, 86077 Pozzilli (IS), Italy*

* *These authors contributed equally*

#### CONSENT

Each participant gave written informed consent before taking part in the study, which was approved by the appropriate local ethical committee at the Institute of Psychiatry, Psychology and Neuroscience, King's College London.

#### DATASET 1

To assess reproducibility and reliability of the proposed methods, diffusion data from 16 healthy adults (8 females) with an age range of 53–65 years was used. All participants were scanned at three time points over a period of one month, having one-week and four-week follow-ups after the first scanning session. The MRI data were acquired using a 3T GE MR750 system equipped with a C-GE HNS 12-channel head coil. For each scanning session, a total of 60 diffusion-weighted MRI volumes (b-value of 1500 $s{\cdot}mm^{-2}$) and nine b=0 $s{\cdot}mm^{-2}$ volumes (including three with reverse phase-encoding direction) were acquired with an isotropic resolution of 2×2×2 $mm^3$ using a spin-echo single shot EPI pulse sequence with echo time TE = 75.4 ms and repetition time TR = 12 s (peripherally pulse-gated). All diffusion data were corrected for artefacts caused by head motion, eddy currents and magnetic field susceptibility distortions using FSL's *eddy* [1] and *topup* pre-processing tools [2,3]. Diffusion tensor and spherical deconvolution model estimation, as well as tractography, were performed using StarTrack (www.mr-startrack.com).

#### DATASET 2

To verify MegaTrack's ability to detect biological changes, MRI data from participants with a diagnosis of motor neurone disease (MND) were compared to data from control participants using conventional manual dissection and MegaTrack. MND provides a well-known example of white matter alteration and results are expected to be localised along the corticospinal tract (CST). Diffusion MRI data were acquired in 25 limb-onset MND patients (51.7 years ± 10.5 years) and 25 controls who were age and gender-matched using a 3T GE HDx Scanner (General Electric, Milwaukee, WI, USA) equipped with an 8-channel head coil. A spin-echo twice refocused EPI sequence was used with a voxel size of 2.4×2.4×2.4 $mm^3$, matrix of 128×128, and a FOV of 307×307 $mm^2$. Overall, 60 contiguous near-axial slices were acquired with TE of 104.5 ms along 32 diffusion-weighted directions with a b-value of 1350 $s{\cdot}mm^{-2}$ and four b=0 volumes. The acquisition was gated to the cardiac cycle using a digital pulse oximeter placed on participants' forefinger, yielding an average TR equal to 14 R-R intervals. The data were pre-processed for motion and eddy current distortion correction using ExploreDTI [4] with the corresponding reorientation of the b-matrix [5]. The diffusion tensor estimation was performed using a non-linear least square approach. Spherical deconvolution estimation was performed using StarTrack.

### DATASET 3

Diffusion MRI data from a cross-sectional adult cohort of 140 participants (68 males, 72 females), aged 18-80 years, were used to build white matter atlases. Diffusion-weighted images were acquired on a 3 T GE Signa HDx TwinSpeed scanner (General Electric, Milwaukee, WI, USA) with an 8-channel head coil. A spin-echo single-shot EPI sequence was used with a voxel size of $2.4\times2.4\times2.4$ mm$^3$, matrix of $128\times128$, and a FOV of $307\times307$ mm$^2$. Overall, 60 contiguous near-axial slices were acquired with an echo time of 93.4 ms along 60 diffusion-weighted directions. A b-value of 3000 s·mm$^{-2}$ was used and seven volumes with no diffusion weighting (b=0) were also collected. The data were acquired using an ASSET factor of 2 and cardiac gating with an effective TR of 20/30 R–R intervals. Datasets were pre-processed for motion and eddy current distortions in ExploreDTI [4] using the T1w image as reference and with the corresponding reorientation of the b-matrix [5]. Model estimation and tractography using the diffusion tensor and spherical deconvolution were performed using StarTrack.
